# Pressure-driven membrane inflation through nanopores on the cell wall

**DOI:** 10.1101/2023.02.19.529118

**Authors:** Qi Zhong, Chen-Xu Wu, Rui Ma

**Affiliations:** Department of Physics, College of Physical Science and Technology, Xiamen University, Xiamen 361005, China; Fujian Provincial Key Lab for Soft Functional Materials Research, Research Institute for Biomimetics and Soft Matter, Xiamen University, Xiamen 361005, China

## Abstract

Walled cells, such as plants and fungi, compose an important part of the model systems in biology. The cell wall primarily functions to prevent the cell from over-expansion when exposed to water, and is a porous material distributed with nanosized pores on it. In this paper, we study the deformation of a membrane patch by an osmotic pressure through a nanopore on the cell wall. We find that there exists a critical pore size beyond which the membrane cannot stand against the pressure and would inflate out through the pore. The critical pore size scales exponentially with the membrane tension and the spontaneous curvature, and exhibits a power law dependence on the osmotic pressure. Our results also show that the liquid membrane expansion by pressure is mechanically different from the solid balloon expansion, and predict that the bending rigidity of the membrane in walled cells should be larger than that of the mammalian cells so as to prevent inflation through the pores on the cell wall.

## Introduction

The cell wall is a structure that surrounds the cells of many organisms, including plants, fungi, bacteria, and some protists [1–4]. It is a rigid layer that provides support, protection, and shape to the cell. In plants, the cell wall is composed primarily of cellulose, a polysaccharide made up of glucose units [5]. The cell wall of fungi is composed of chitin, a polymer of N-acetylglucosamine [6, 7]. Bacterial cell walls are made up of peptidoglycan, a polymer of amino sugars and amino acids [8, 9]. The cell wall plays an important role in maintaining the integrity of the cell and protecting it from mechanical stress, osmotic pressure, and other environmental factors.

Basically, the cell wall is a porous structure that allows for the exchange of materials between the cell and its environment. The size and shape of the pores in the cell wall can vary depending on the type of cell and the function of the wall. For example, in the primary cell wall of plant cells, the size of the pores can range from 10-20 nanometers to several micrometers in diameter, with the largest pores often found at the corners where adjacent cells meet [10, 11]. In fungal cell walls, the size of the pores can also vary, but is typically at an order of a few tens of nanometers [12, 13].

Walled cells generally have a high osmotic pressure that allows them to grow and survive in the environment [14–19]. The osmotic pressure pushes the plasma membrane against the cell wall and therefore prevents some important biological process from happening. In particular, endocytosis which involves the internalization of a small patch of plasma membrane is hindered by the osmotic pressure [20]. In fission yeast, the turgor pressure is estimated to be 0.85 ± 0.15 MPa by measuring the growth curve of a mutant yeast cell against a PDMS chamber [21, 22]. In another experiment which compares the geometry of a fission yeast cell upon osmolarity change, it is estimated that the turgor pressure is up to 1.5 ± 0.2 MPa [14]. Based on a similar osmolarity variation method, the value of turgor pressure is estimated to be 0.6 ± 0.2 MPa in budding yeast [23]. In general, the turgor pressure in yeast cells have an order of magnitude 0.1 – 1 MPa.

In this work, we study membrane deformation through a nanopore on the cell wall by the osmotic pressure with the Helfrich model. We find that when the pore is small, the membrane assumes a certain shape to resist the osmotic pressure. When gradually increasing the pore size beyond a critical value, the membrane cannot stand against the pressure anymore and would inflate out through the pore and further expand. We systematically study how the critical pore size *R*_crit_ depends on the membrane properties and find that *R*_crit_ scales exponentially with the membrane tension and the spontaneous curvature, but scales with the osmotic pressure with a power law. We show that the expansion of a liquid membrane by pressure is different from a solid one in terms of their *p-V* curve. Our results suggest that the membrane bending rigidity in walled cells should be larger than that in mammalian cells so as to prevent the cell from inflation through the nanopores present on the cell wall.

## Methods

### Model

We consider a walled cell with a high osmotic pressure *p* that pushes the cell membrane against the cell wall. A nanopore with a radius of *R*_pore_ is present on the cell wall, which has a much larger Young’s modulus than the osmotic pressure and a much smaller curvature than the nanopore [24–27]. The cell wall is therefore modeled as a rigid and flat substrate. We assume the deformation of the membrane at the pore is axisymmetric, and parameterized with its meridional coordinates [*r*(*u*), *z*(*u*)], where *u* ∈ [0, 1] is the rescaled arclength. The point at *u* = 0 labels the membrane tip and the points at *u* =1 label the membrane edge (Fig. 1b). The coordinates satisfy the geometric relations

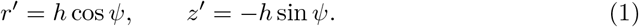

Hereafter, we use *f’* to denote the derivative of an arbitrary function f with respect to *u*. The angle *ψ*(*u*) spans between the tangential direction and the horizontal direction. The constant *h* is essentially the total arclength of the membrane profile and is to be solved with the shape equations. The total energy of the membrane reads

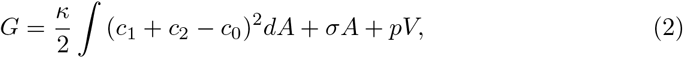

where the first term is the classical Helfrich bending energy with *κ* denoting the bending rigidity of the membrane, 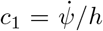 and *c*_2_ = sin *ψ/r* the two principal curvatures of the membrane surface, *c*_0_ denoting the spontaneous curvature induced by curvature-generating proteins coated on the membrane [28, 29]. In the absence of proteins, the spontaneous curvature is assumed to vanish. The second term in Eq. (2) is the surface tension energy with *A* denoting the surface area of the deformed membrane, and *σ* denoting the membrane tension at the edge of the pore. The third term describes the effect of the osmotic pressure with *V* being the increased volume of the membrane due to inflation, and *p* denoting the osmotic pressure (Fig. 1a). Due to the rotational symmetry of the membrane shape about the *z* axis, the total energy of the membrane in Eq. (2) can be expressed as a functional of the shape variables

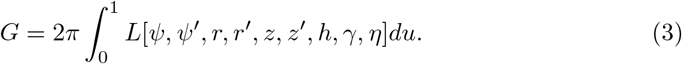

Here *γ, η* are Lagrangian multipliers that enforce the geometric relations in Eq. (1). The shape equations of the membrane are obtained by applying variations of the free energy *G* with respect to all the shape variables [30, 31]. The variation *δG* contains both bulk terms, e.g., [*∂L/∂ψ* – *d*(*∂L/∂ψ’*)/*du*] *δψ*, and boundary terms, e.g., 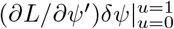. We obtain the shape equations by having the bulk terms to be zero. In addition, proper boundary conditions (BCs) at the membrane tip *u* = 0 and at the pore edge *u* =1 need to be specified, which can be achieved by setting the boundary terms in *δG* to be zero. There are two types of BCs, the free-hinge BC in which the membrane angle *ψ* is allowed to freely rotate, and the fixed-hinge BC in which the membrane angle *ψ* is fixed to be 0 (Fig. 1b). They correspond to have either *∂L/∂ψ’* = 0 or *δψ* = 0 in the boundary term. The detailed derivation of the shape equations and the BCs is provided in the appendix.

We numerically solve the shape equations with the bvp4c solver in MATLAB, which is designed for solving boundary value problems (BVPs) of ordinary differential equations.

**Figure 1.**
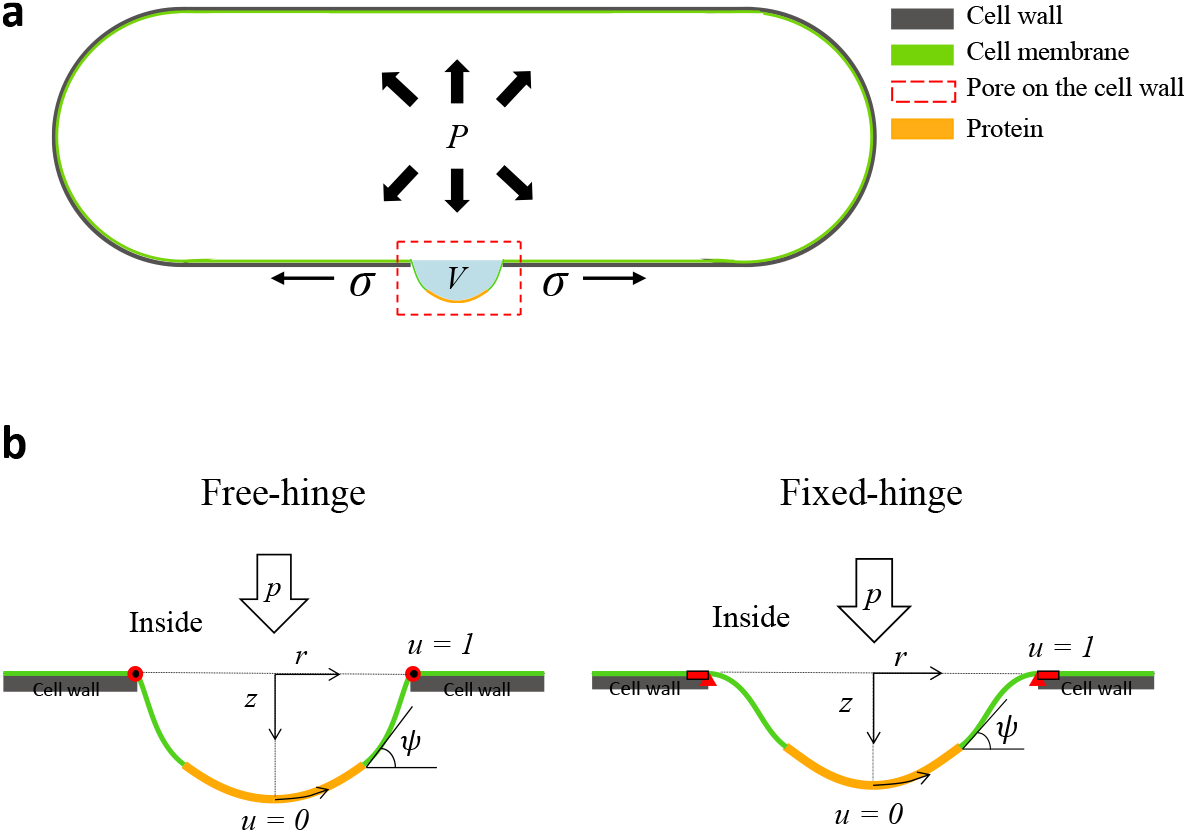
Schematic illustration of a typical yeast cell and the model. **(a)** Illustration of the typical morphology of a fission yeast cell. A nanopore (red frame) is present on the cell wall (gray) and the membrane (green) is pushed outwards by the osmotic pressure *p*. The size of the nanopore is exaggerated for the purpose of illustration only. **(b)** Illustration of the membrane models. The membrane shape is assumed to be axisymmetric with respect to the z-axis, and parameterized by its meridinal coordinates [*r*(*u*), *z*(*u*)] with *u* = 0 labeling the membrane tip and *u* =1 labeling the membrane edge. Two types of boundary conditions (BCs) are considered, namely the free-hinge BC (left) in which the membrane is allowed to freely rotate at the pore edge and the fixed-hinge BC (right) in which the membrane angle is fixed. The proteins (orange) coated on the membrane are assumed to generate the spontaneous curvature of the membrane.

## Results

### Critical pore size for membrane inflation scales exponentially with the membrane tension and spontaneous curvature

In this section, we fix the osmotic pressure *p* and study how the membrane is deformed when gradually increasing the pore radius *R*_pore_. The osmotic pressure *p* and the membrane bending rigidity *κ* define a characteristic length *R_p_* = (*κ*/2*p*)^1/3^, which is used to rescale the length, and nondimensionalize all the other parameters. In particular, the rescaled pressure 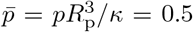 is a constant. The other nondimensionalized parameters include the rescaled tension 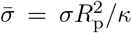, and the rescaled spontaneous curvature 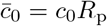.

We first consider a membrane without any curvature-generating proteins and study how the membrane is deformed when continuously increasing the radius of the pore *R*_pore_ under a constant membrane tension 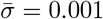. The membrane is pushed down by the pressure *p* to a depth of *D*_pore_. It is found in the *D*_pore_ – *R*_pore_ curve, a single value of the pore radius *R*_pore_ corresponds to two membrane shapes (squares and circles in Fig. 2 c), one with a shallow depth and the other with a deep one. By comparing the total energy of the two shapes (Fig. 2 b), we see that the energy of the shallow solution is much lower than that of the deep one, indicating an energetically more favorable state of the membrane. For the shallow shapes, the depth of the membrane is pushed further down by the pressure as the pore size is enlarged, while for the deep shapes, the depth of the membrane shows a nonmonotonic relation with the pore radius. There exists a critical pore size *R*_crit_ beyond which no solutions are found, which indicates that the osmotic pressure is too strong for any membrane shape to sustain the pressure. The membrane would continuously inflate out of the pore and expand over time if there is constant supply of lipid molecules flowing into the pore from the edge.

**Figure 2.**
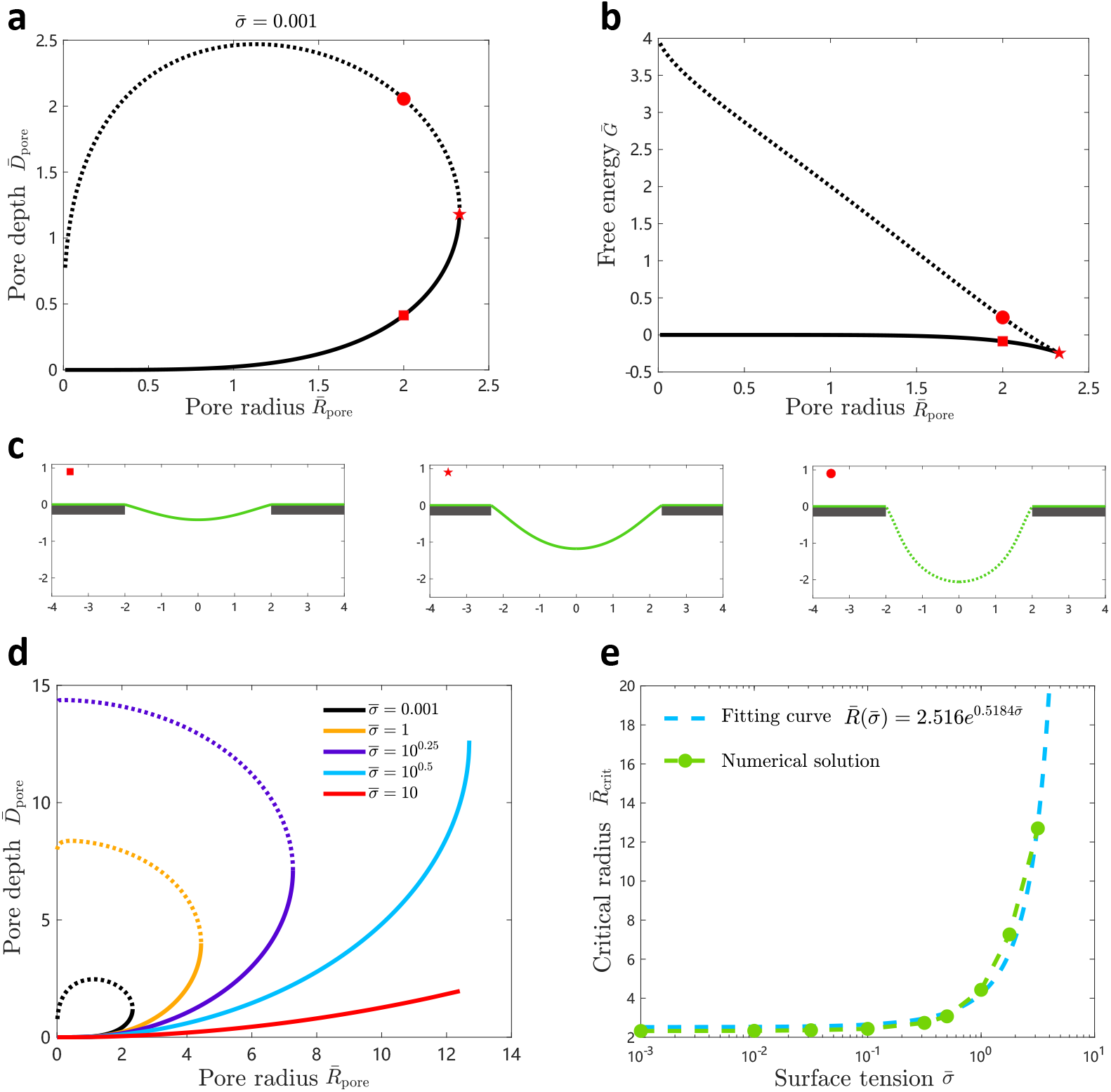
Effect of the surface tension on the critical radius of an uncoated membrane. **(a)** *D_pore_* vs. *R*_pore_ curve of membrane deformations for an uncoated membrane. The critical pore size, beyond which no solutions are found, is indicated by a pentagon. **(b)** The free energy curve of the corresponding membrane deformations in (a). The solid curve indicates a stable state, and the dashed line indicates a metastable state. **(c)** Illustration of the membrane shapes indicated by the corresponding symbols on the *D*_pore_ vs. *R*_pore_ curve. **(d)** *D*_pore_ vs. *R*_pore_ curves are shown for different membrane tensions 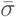. **(e)** The critical pore size as a function of the surface tension 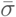. The green dotted curve represents the numerical solutions. The cyan dashed curve represents the exponential fitting of the numerical solution.

We then plot the *D*_pore_ vs. *R*_pore_ curves for different membrane tensions 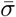 (Fig. 2)d. The critical pore size 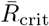 is found to strongly depend on the membrane tension 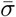. For 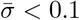, the critical radius 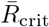 stays around 2. Beyond 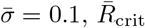 has a dramatic increase with 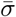 (Fig. 2e, green dotted curve). The relationship can be nicely fitted with an exponential function (Fig. 2 e, cyan dashed curve).

We next consider a membrane totally coated by proteins and study the effect of spon-taneous curvature c0 induced by the proteins on the membrane deformation, particularly on the critical pore radius. The *D*_pore_ – *R*_pore_ curve for a coated membrane with positive *c*_0_ show a similar trend with that of the uncoated membrane, i.e., a single pore radius corresponding to two solutions, one with a lower energy and the other with a higher one (Fig. 3a, b). However, solutions with negative depth appear for small pore radii in the lower energy branch due to the positive spontaneous curvature c0 that tends to bend the membrane inward against the osmotic pressure (Fig. 3a). When the pore radius becomes large, the spontaneous curvature *c*_0_ cannot resist the osmotic pressure anymore and the membrane is pushed outward by the pressure (Fig. 3c). The *D*_pore_ – *R*_pore_ curves for different spontaneous curvatures *c*_0_ are shown in Fig. 3d. The critical radius increases with the spontaneous curvature and the relationship can be well fitted by an exponential function (Fig. 3e).

**Figure 3.**
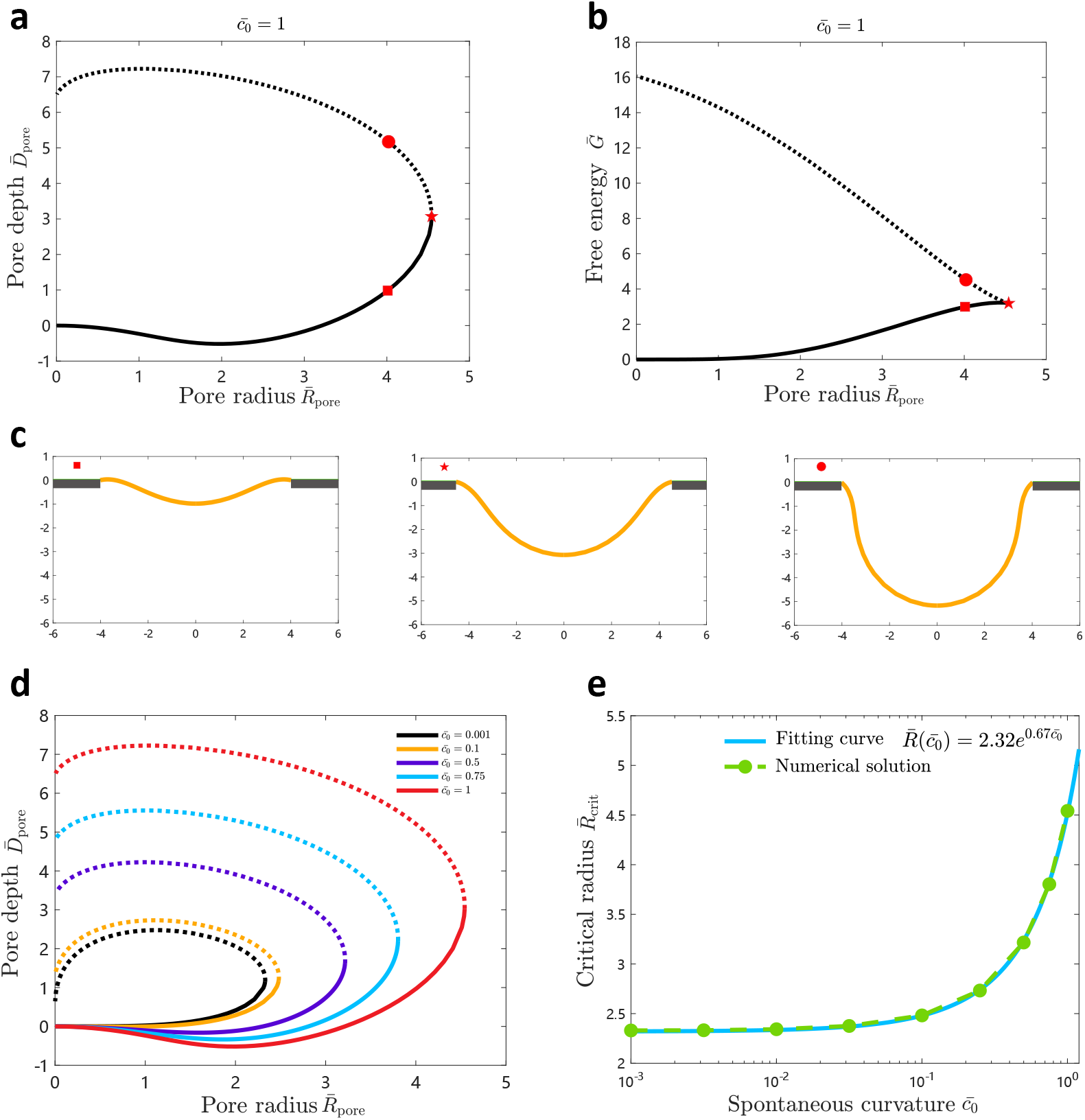
Effect of the spontaneous curvature on the critical radius of a totally coated membrane. **(a)** *D_pore_* vs. *R_pore_* curve of membrane deformations for a totally coated membrane with spontaneous curvature 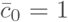. The critical pore size is indicated by a pentagon. **(b)** The free energy curve of the corresponding membrane deformations in (a). The solid line indicates a stable state, and the dashed line indicates a metastable state. **(c)** Illustration of the membrane shapes indicated by the corresponding symbols on the *D*_pore_ vs. *R*_pore_ curve. **(d)** *D*_pore_ vs. *R*_pore_ curves are shown for different spontaneous curvatures 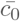. **(e)** The critical pore size as a function of the spontaneous curvature 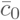. The green dotted curve represents the numerical solutions. The cyan dashed curve represents the exponential fitting of the numerical solution.

### Critical pressure for membrane inflation scales with the pore radius with a power law

In this section, we fix the pore radius and study how an uncoated membrane is deformed by gradually increasing the osmotic pressure *p*. The membrane tension *σ* and the membrane bending rigidity *κ* defines a characteristic length *R_σ_* = [*κ*/(2*σ*)]^1/2^, which is used to rescale the length, and nondimensionalize all the other parameters. In particular, the membrane tension can be rescaled to a constant 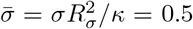. The rescaled osmotic pressure is 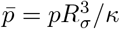.

A typical 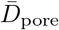 vs. 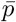 curve is shown in Fig. 4a. A single osmotic pressure 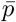 can find two corresponding depths on the curve (Fig. 4), with the shallow one having a lower energy and the deep one having a higher energy (Fig. 4b, c). We stress that for small osmotic pressure p, the high energy branch can have a very large depth. This is similar to the *p-V* curve of an inflating balloon [32–36]. When the osmotic pressure is a beyond a critical value, no solutions are found. It implies that the osmotic pressure is too large for any membrane shape to sustain the pressure. The membrane would inflate out of the pore to form a bulge. The critical pressure 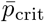 is found to increase with the pore radius (Fig. 4d), and the relationship can be well fitted with a power law 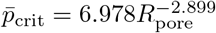 (Fig. 4e).

**Figure 4.**
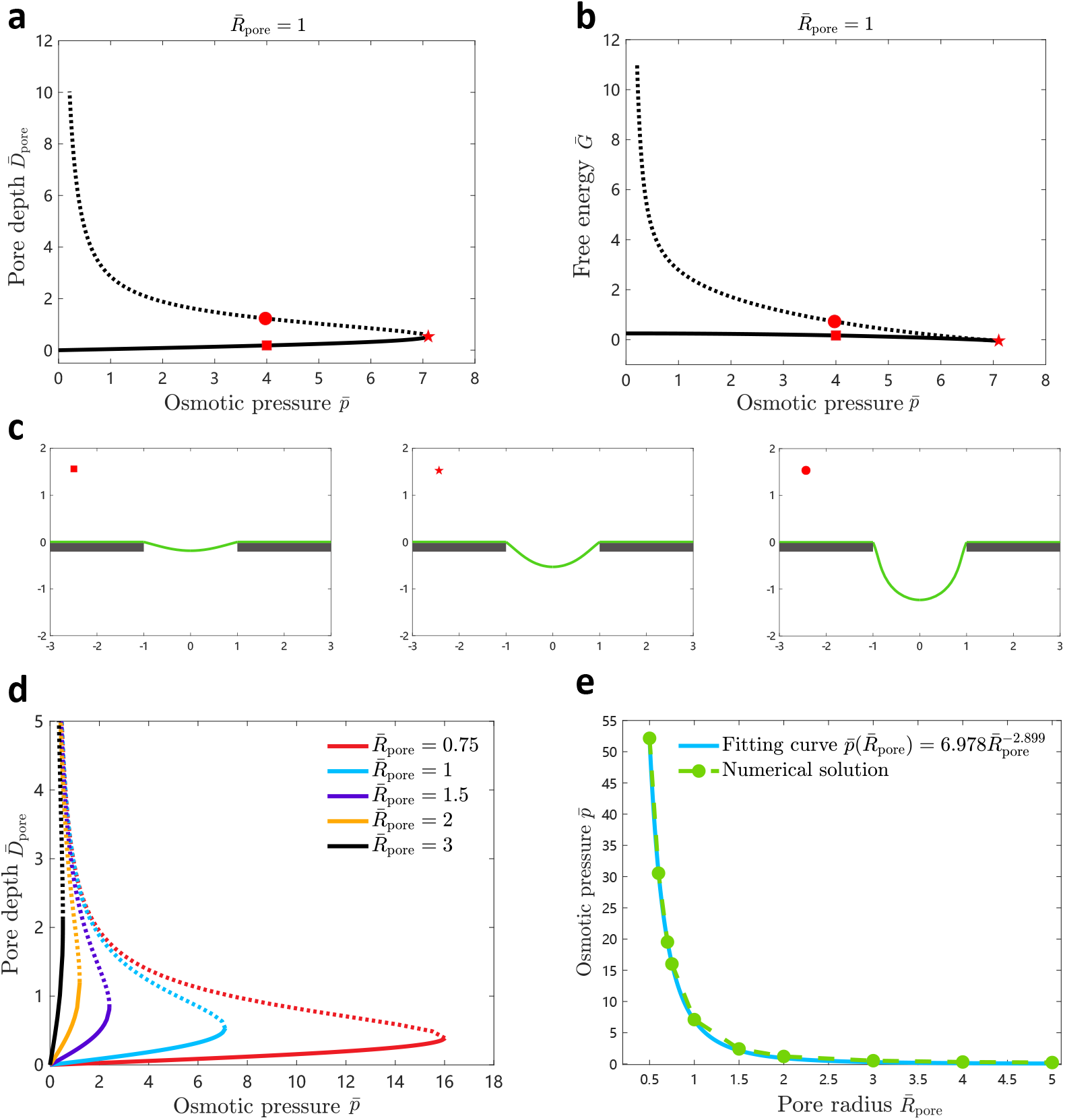
Effect of the osmotic pressure on the membrane deformation for an uncoated membrane. **(a)** *D*_pore_ vs. 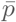 curve of membrane deformations for an uncoated membrane. The pore size is fixed at 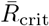 = 1. The critical osmotic pressure is indicated by a pentagon. **(b)** The free energy curve of the corresponding membrane deformations in (a). The solid curve indicates a stable state and the dashed curve indicates a metastable state. **(c)** Illustration of the membrane shapes indicated by the corresponding symbols on the *D*_pore_ vs. 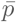 curve. **(d)** *D*_pore_ vs. 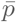 curves are shown for different pore size 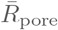. **(e)** The critical pressure as a function of the pore size 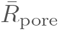. The green dotted curve represents the numerical solutions. The cyan dashed curve represents the power law fitting of the numerical solution.

## Discussion

### The critical pore size is about 10 nm given the typical parameters of the membrane

We have shown that if a nanopore is present on the cell wall and the pore radius is small, the membrane is able to resist the osmotic pressure by assuming a curved shape. If the pore radius is beyond a critical value *R*_crit_, no membrane shapes are able to sustain the osmotic pressure and the membrane would inflate out and further expand. In Fig. 5, we show how the critical pore size depends on the model parameters, given the typical value of the membrane properties. The model parameters include the bending rigidity *κ*, the membrane tension *σ*, the osmotic pressure p, and the spontaneous curvature *c*_0_. We fix three of them to the typical values (listed on the top of each panel in Fig. 5), and vary the fourth one to see how it impacts the critical pore size *R*_crit_ using the fitted expression 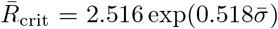 for the free-hinge BC and 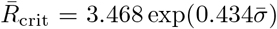 for the fixed-hinge BC. The critical pore size *R*_crit_ is found to decrease with the osmotic pressure and increase with the other three parameters, and stays in the range of 5 – 20 nm. In general, *R*_crit_ is smaller under the free-higne BC than that under the fixed hinge BC. As we have pointed out, the typical pore size of the cell wall is about tens of nanometers [10–13], which is comparable to the critical pore size calculated by our theory (Fig. 5b). Our calculation therefore implies that the bending rigidity of the walled cells must be higher than the value 20 kT we have typically assumed. This is plausible based on the experimental evidence that the membrane is surrounded with actin meshwork in yeast cells [37–39], which is able to effectively increase the bending rigidity of the membrane. Furthermore, proteins on the membrane are also help to increase the bending rigidity.

**Figure 5.**
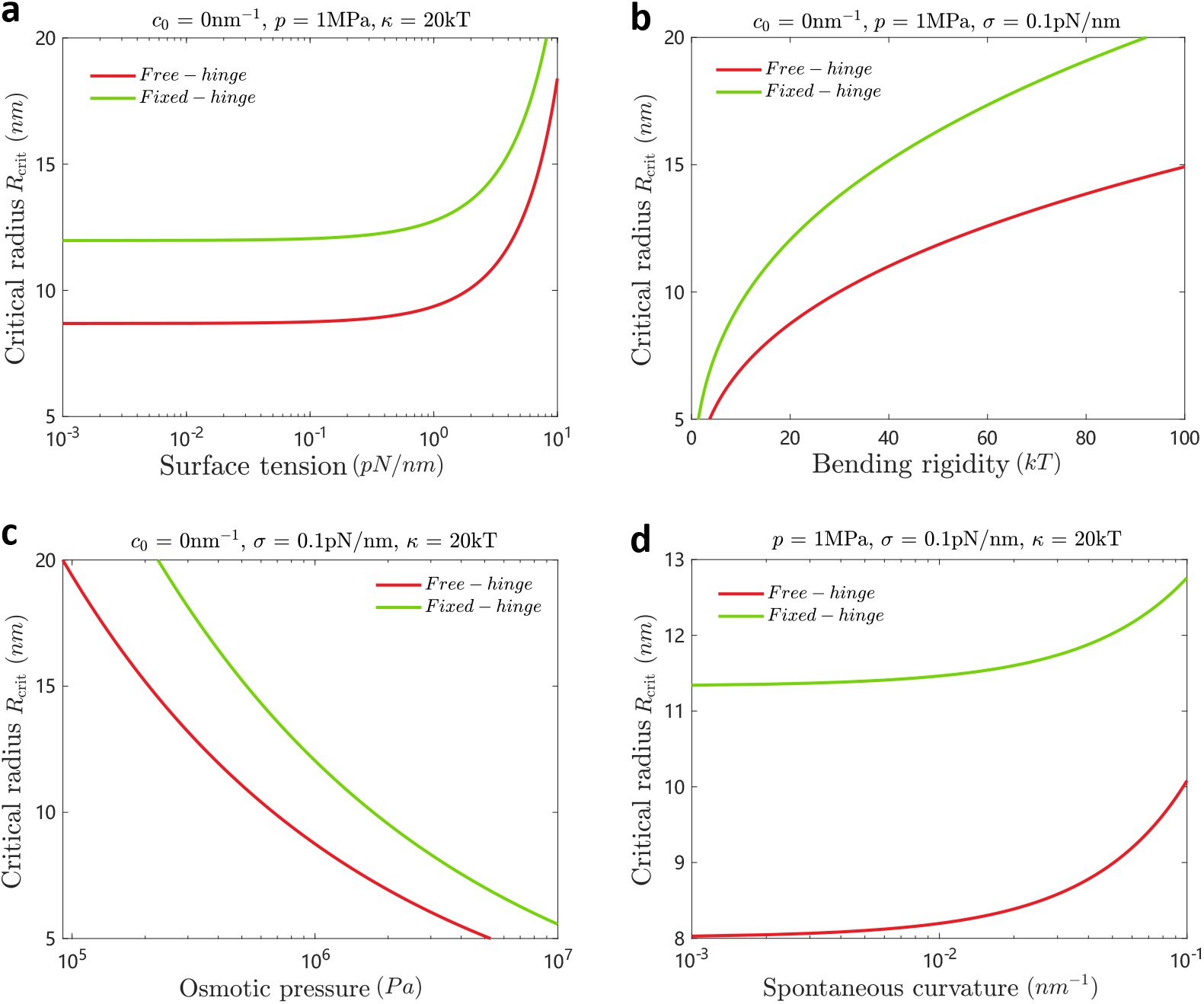
Dependence of the critical pore size on the model parameters. The typical values of the fixed parameters are listed on top of each panel. The red curve is for the free-hinge BC and the green curve is for the fixed-hinge BC. **(a-d)** The relationship between the critical radius *R*_crit_ and the surface tension *σ*, the bending rigidity *κ*, the osmotic pressure *p*, and the spontaneous curvature *c*_0_, respectively.

### Liquid membrane versus solid balloon

In Fig. 4, we have presented the results for the membrane deformation when gradually increasing the osmotic pressure. The *D*_pore_ vs. 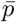 curve is found to be nonmonotic, which resembles the *p-V* curve of an inflating balloon [32–36]. In practice, the setup shown in Fig. 4 is used to measure the elastic properties of solid polymer materials, such as rubber. However, for the solid balloon, in particular, at very large volume, the *p-V* curve would increase with the pressure again due to the stretching energy of the rubber that makes up the balloon. In our case, we consider an incompressible liquid membrane and the membrane area increase is due to the lipid flow from the pore edge when the membrane is pushed by the osmotic pressure. The membrane tension *σ* at the edge is fixed instead of increasing with the depth. Therefore, we can find deep invaginations at very small pressure, which differs the liquid membrane from the solid balloon.

## Conclusion

We have studied membrane inflation through a nanopore on the cell wall in the presence of a high osmotic pressure. It is found that there exists a critical value of the pore size *R*_crit_ beyond which the membrane cannot stand against the osmotic pressure and would inflate out through the pore and keep expansion. The critical pore size *R*_crit_ scales exponentially with the membrane tension *σ* and the spontaneous curvature *c*_0_, and scales with the osmotic pressure *p* with a power law. The boundary conditions at the pore edge also make a quantitative difference. In general, the fixed-hinge BC leads to a larger *R*_crit_ than the free-hinge BC. Our results also reveal that the inflation of a liquid membrane by pressure behaves differently than that of a solid balloon. In walled cells where the osmotic pressure is high, the membrane should have a large bending rigidity than that in mammalian cells to prevent from inflation through the pores on the cell wall.

## Appendix

### Derivation of the membrane shape equations

In order to derive the membrane shape equations, we explicitly express the integrand *L* in Eq. (3) as

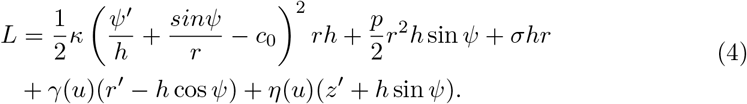

Here the three terms in the first line describe the energy contribution made by the bending rigidity, the surface tension, and the osmotic pressure, respectively, and the two terms in the second line describe the Lagrangian multiplies *γ*(*u*) and *η*(*u*) to enforce the geometric constraints Eq. (1). The variation of the functional *G* in Eq. (3) reads

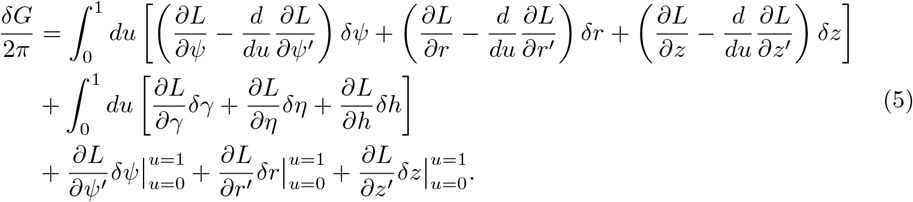

The variation contains both the bulk terms (first and second lines) and the boundary terms (third line). By letting the bulk terms equal to zero, we obtain a set of ordinary differential equations, which include a second-order equation

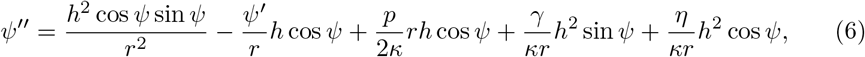

and two first-order equations

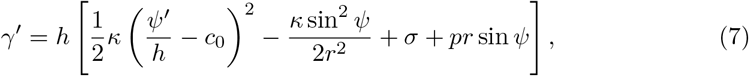

and

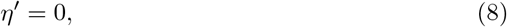

as well as the two geometric relations in Eq. (1).

In addition, the equation *∂L/∂h* = 0 does not give an equation of *h*, but leads to a conserved quantity [30, 31]

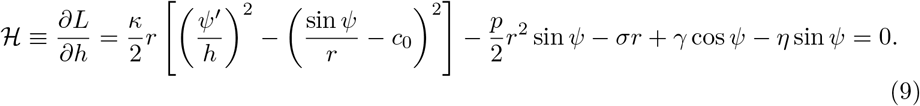

There is a freedom to choose the scale function *h*. Here we choose *h’* = 0 such that *h* is a constant to be determined. It is essentially the total arclength of the membrane profile.

### Derivation of the boundary conditions

By letting the boundary terms in Eq. (5) equal to zero, we obtain the BCs. In particular, in the boundary term about *δr*, we let 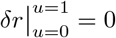, which implies to fix the radius

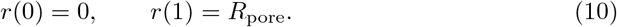

In the boundary term about *δz*, we obtain

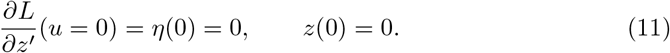

As for the boundary term about *δψ*, at the membrane tip, we have

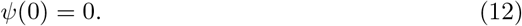

At the pore edge, we have two choices. If we let *δψ*(1) = 0, it is equivalent to the fixed-hinge BC

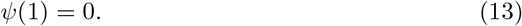

If we let *∂L/∂ψ’* = 0, it is equivalent to the free-hinge BC

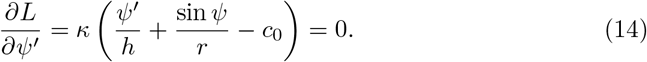

The conserved quantity 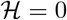 at the membrane tip *u* = 0 gives rise to a BC

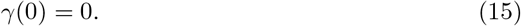

### Numerical methods to calculate the *D_pore_* vs. *R_pore_* curve

So far we have obtained a total number of 5 equations, which include Eqs. (1), (6), (8), (8). Since only Eq. (6) is first order, they are equivalent to a total number of 6 first order ordinary differential equations. In addition, Eqs. (10), (11), (12), (13) or (14) constitute a total number of 7 BCs. Overall, the 6 equations plus 1 unknown parameter *h*, and the 7 BCs constitute a well-defined boundary value problem.

We solve the problem with the matlab solver bvp5c, which requires the equations and the BCs as its input. Furthermore, the solver requires an initial guess of the solutions. In order to obtain the *D*_pore_ vs. *R*_pore_ curve, We always start with a small *R*_pore_ and use the flat shape as our initial guess. In this case, the membrane is almost flat, and it is easy for the solver to find the solution. We then gradually increase the pore radius *R*_pore_ to *R*_pore_ + ΔR and use the solution for *R*_pore_ as the initial guess to solve the equation for *R*_pore_ + ΔR. This iteration step allows us to extend the solution to the critical pore radius since beyond which no solutions are found anymore. We then use the solution at the critical pore radius as the starting point and iterate over the membrane depth *D*_pore_ to obtain the other branch of the solution. During this iteration, we add the boundary condition *z*(0) = *D*_pore_ and set the pore radius *R*_pore_ as an unknown parameter to be solved by the solver.

## Acknowledgments

We acknowledge financial support from National Natural Science Foundation of China under Grants No. 12004317, Fundamental Research Funds for Central Universities of China under Grant No. 20720200072, and 111 project No. B16029.

